# FFixR: A Machine Learning Framework for Accurate Somatic Mutation Calling from FFPE RNA-Seq Data in Cancer

**DOI:** 10.1101/2025.10.17.682574

**Authors:** Or Livne, Keren Yizhak

**Affiliations:** Department of Cell Biology and Cancer Science, The Ruth and Bruce Rappaport Faculty of Medicine, Technion - Israel Institute of Technology, Haifa 3525422, Israel

## Abstract

Formalin-fixed paraffin-embedded (FFPE) tissues are widely used in clinical and research settings, yet their use for detecting somatic mutations from RNA sequencing (RNA-seq) is hindered by artefactual mutations introduced by cytosine deamination and strand-specific damage. Existing FFPE noise-filtering tools are tailored to DNA-seq and rely on strand bias, rendering them unsuitable for RNA-seq. Here, we present FFixR, a machine learning–based framework that filters FFPE-induced artefacts from RNA-seq data without requiring matched-normal samples. Trained on FFPE melanoma samples with matched DNA, FFixR leverages allele-specific read counts, variant features, and mutational signature probabilities. FFixR removed up to 98% of artefactual mutations while maintaining ∼92% recall of true variants. SHAP analysis revealed key feature interactions guiding model decisions. When applied to an independent cohort, FFixR restored the correlation between RNA- and DNA-derived tumor mutational burden (R² = 0.881) and recovered biologically meaningful mutational signatures. FFixR enables accurate somatic variant calling from FFPE RNA-seq data, expanding the utility of archival samples for research and clinical applications.

## Background

Formalin fixation and paraffin embedding (FFPE) is a common technique used in clinical tissue preservation. In contrast with freeze preservation, this method ensures long-term storage at room temperature, which allows for a much simpler tissue collection and storage. While FFPE-preserved specimens are used in various diagnostic applications, this preservation introduces noise into the analysis of DNA and RNA sequencing that needs to be properly addressed. Specifically, formalin fixation leads to the fragmentation of the genetic material and subsequently to hydrolytic deamination of cytosine^1–3^. In practice, this process replaces cytosine with deoxyuridine, which leads to the formation of C:G>T:A or C:G>U:A mutations^1–3^ in DNA or RNA, respectively. Following, multiple artefacts are detected when searching for somatic point mutations in DNA or RNA sequencing data. In fact, the amount of FFPE induced mutations can be two orders of magnitude more than the true mutations, even in highly mutated tumors, raising the need for detecting and eliminating this source of noise.

In recent years several computational tools were developed in order to tackle this problem and filter the FFPE noise. All of these tools utilize a feature that describes the mutation’s strand bias as their key feature. True somatic mutations are expected to be represented on both DNA strands, as a consequence of cellular replication and DNA correction processes in living tissue. In contrast, FFPE-induced artefacts occur after cellular death, and therefore typically manifest on a single strand without being propagated to its complement. For example, FilterByOrientationBias (GATK)^4^ filters Mutect2 somatic variant calls based on the assumption that artifacts are generally strand biased. SOBDetector^5^ is mainly based on alternative allele’s orientation and number of reads and fraction. FFPolish^6^ is based on allele frequency, size of insert fragment and strand bias. DEEPOMICS FFPE^7^ is a neural network tool, based mainly on alternate and reference alleles’ frequency and read depth, mutation sequence (e.g. C>T), mutation type (deletion, insertion, etc.) and strand orientation bias. In their publication, Heo et al.^7^ evaluated the performance of exiting FFPE noise-filtering tools on their own dataset, reporting moderate to high recall (0.681-0.969) but very low precision (0.024-0.129) for methods such as FilterByOrientationBias^4^, SOBDetector^5^ and FFPolish^6^. In contrast, their own model achieved improved overall performance on the same dataset, yielding an MCC of 0.721, precision of 0.742, and recall of 0.708.

In recent years RNA sequencing data has proven useful in detecting somatic mutations in different contexts, including for studying somatic clonality in normal tissues and for estimating tumor mutational burden in cancer samples^8–10^. However, identifying somatic mutations from FFPE RNA-seq data remains a challenge. Specifically, since RNA is single-stranded, the strand bias feature cannot be utilized, making previously published tools designed for DNA incompatible for this type of data.

In this study, we developed an ML-based framework for filtering FFPE-induced artifacts in RNA-seq data. The approach is based on a machine learning model trained on features derived from RNA-seq data, including alternative allele count and frequency, predicted somatic probability, variant classification, sequence context and cosine similarity to mutational signatures^11^. Additionally, we applied mathematical transformations to selected features to enhance the model’s ability to discriminate between true variants and noise. The resulting model effectively removed the majority of FFPE-induced artifacts while preserving high sensitivity for true variant detection, underscoring its utility in both clinical diagnostics and research-oriented next generation sequencing applications.

## Results

### Eliminating FFPE originated sequencing noise in RNA-seq data

To filter FFPE artefacts from bulk RNA-seq data we leveraged FFPE-preserved DNA and matched RNA sequencing data from 39 melanoma patients available in the Van Allen et al. dataset^12^. As our ground truth we used the set of point somatic mutations detected in DNA sequencing data following FFPE filtering. To obtain the initial set of RNA-based candidate somatic mutations, we applied our previously developed RNA-MuTect-WMN method^8^ which detects somatic mutations from RNA-seq without a matched-normal sample and applies a set of RNA-specific filtering criteria^9^.

To classify the extracted point mutations as true (detected in the DNA) or noise (not detected in DNA and potentially FFPE artefacts), we collected for each variant a set of features (Supp Table 1). Those include the alternate read counts, the mutations’ allele frequency, the transcript strand and variant type. In addition, we incorporated scores computed by MuTect that estimate the probability of a given mutation being a true somatic event rather than sequencing noise. Finally, we included sequenced-based features such as the mutation’s tri-nucleotide context and the probability of association with mutational signatures, as derived from the SignatureAnalyzer tool^13^ (Methods). Specifically, four signatures were detected in this data. Signatures 7 and 11 which are common in melanoma, the common signature 1 that is mostly associated with aging and signature 30 with an unknown etiology. To make our model generic rather than melanoma-specific, we included only signatures 1 and 30.

To evaluate the individual predictive value of each feature in distinguishing true mutations from FFPE-induced artifacts, we assessed their standalone performance using precision, recall, and Matthews correlation coefficient (MCC) as evaluation metrics. MCC is particularly suitable for imbalanced datasets like ours, and served as our primary optimization criterion. We found that the overall performance of each feature alone is low, as demonstrated by the low precision values (Fig. 1a). These results emphasize the need for a more sophisticated model that can aggregate and weight different features to receive an optimal classification.

**Fig 1.**
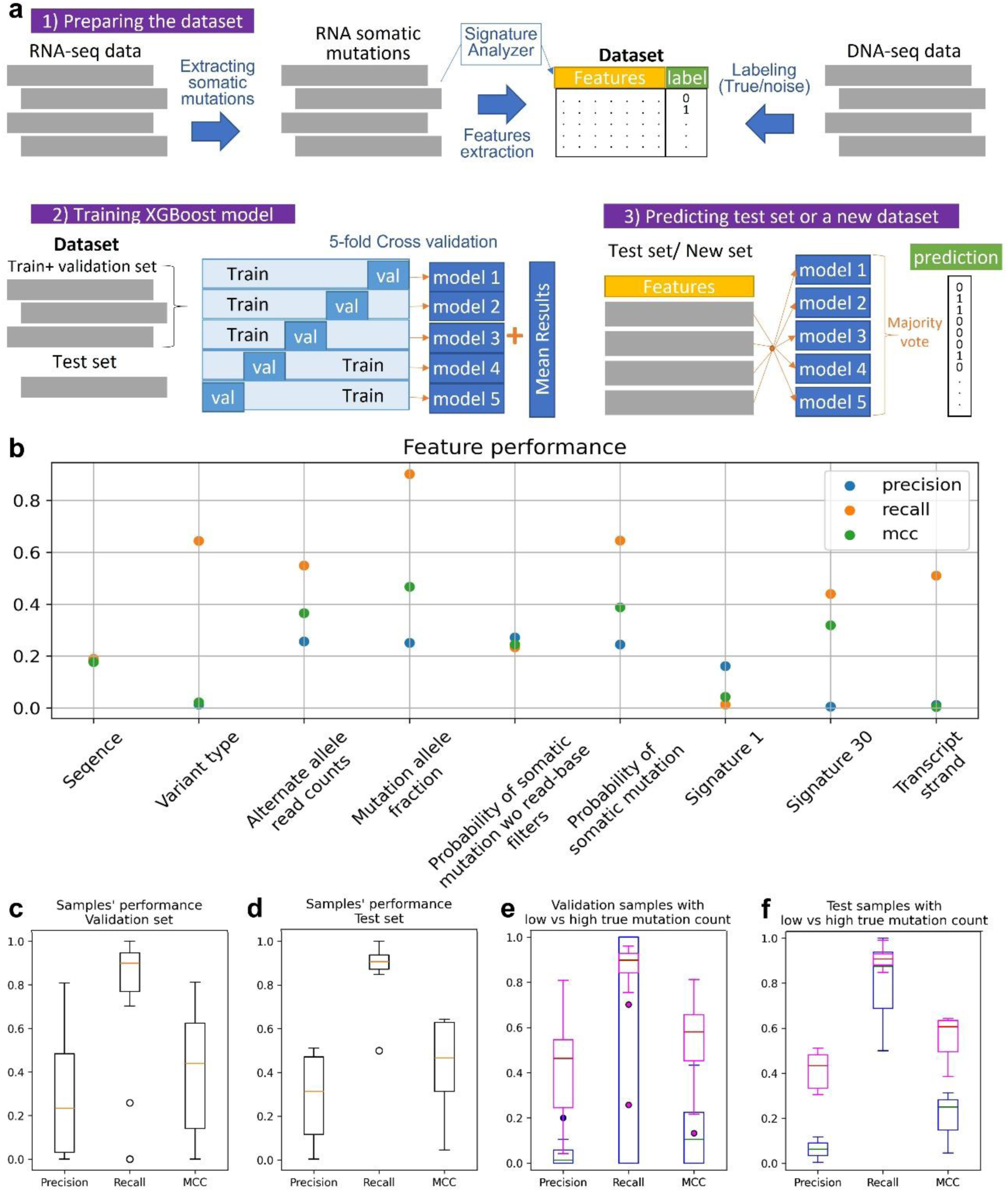
Summary of pipeline predictions. **a** An overview of the pipeline: 1) Data preparation includes the extraction of candidate somatic mutations and their features from RNA-seq data, and labeling mutations as either true or noise based on matched DNA sequencing data 2) For model training we split the dataset into train-validation sets and test set, training XGBoost models in a 5-fold cross validation manner. 3) For model testing we classified each mutation in the left out test set or an external dataset, based on majority vote of the 5 trained models. **b** Prediction power of each feature on its own, considering the optimal threshold. **c-d** Distribution of precision, recall and MCC values on the validation (c) and test (d) sets. Box plots show median, 25th, and 75th percentiles. The whiskers extend to the most extreme data points not considered outliers, and the outliers are represented as dots. **e-f** Distribution of precision, recall and MCC in samples with high (>20) and low (≤20) mutation count, in the validation (e) and test (f) sets.

To this end, we developed a machine learning framework, designated **FFixR** (**F**ormalin-**Fix**ed **R**NA-seq artefact remover), designed to classify variants as either true mutations or FFPE-induced artifacts, based on their associated feature profiles. The model was trained using eXtreme Gradient Boosting (XGBoost), a robust ensemble learning method. For training and validation, we randomly selected 30 samples from the dataset and implemented a 5-fold cross-validation strategy. Each fold produced an independent model that was then used to evaluate the performance in the held-out validation set. Classification of variants in the test set, comprising the remaining 9 held-out samples, was performed using majority voting across the 5 models (Fig. 1b). This ensemble approach was employed to enhance robustness and generalizability of the classifier.

Across the validation sets, a total of 188,075 mutations labeled as noise were evaluated, of which 180,624 (96%) were correctly identified and excluded by the model. Only 576 out of 3,950 true mutations were misclassified, corresponding to a recall of 0.854, precision of 0.312, and a Matthews Correlation Coefficient (MCC) of 0.501 (Fig. 1c). On the held-out test set, the model demonstrated slightly improved performance, eliminating 72,054 out of 73,851 noise mutations (97.6%) while incorrectly discarding only 63 of 824 true mutations. This yielded an MCC of 0.516, precision of 0.297, and recall of 0.924 (Fig. 1d). Further stratified analysis revealed that samples containing a low number of true mutations (≤20) exhibited markedly lower precision and MCC compared to those with higher mutation counts (>20), indicating performance is sensitive to class distribution within individual samples (Fig. 1e,f).

We next compared the performance of FFixR to that achieved by Sorkin et al.^10^ which also introduced a pipeline for filtering FFPE noise mutations from RNA-seq data. While their model also employed an XGBoost classifier with a partially overlapping feature set, a major difference is their reliance on DNA-oriented tools like Mutect2, whereas our approach utilizes RNA-MuTect-WMN^8^, a pipeline specifically optimized for RNA-seq data. Indeed, applying their pipeline to our dataset yielded only 7,658 true mutations out of 4,545,928 total calls (∼0.2%). Moreover, when implementing their model for FFPE noise filtering it labeled all mutations as noise, both true and false mutations, thus achieving a recall and precision of 0, and MCC of −0.041.

Finally, we benchmarked XGBoost against several alternative models, including a neural network, logistic regression, random forest, and LightGBM. XGBoost consistently outperformed the others in terms of MCC. The neural network, in particular, required significant training time and performed poorly unless the training data were balanced.

### Explaining precision performance by analyzing false positive mutations

Precision-recall analysis across varying classification thresholds revealed an atypical pattern (Fig. 2a). Conventionally, increasing the classification threshold leads to a reduction in recall—due to the exclusion of more true positives—while typically improving precision, as false positives are eliminated at a faster rate than true positives. However, in our model, increasing the threshold did not yield a corresponding improvement in precision. This suggests that the misclassified false positives exhibit similar feature profiles to the true positives, making them difficult to distinguish. To investigate this further, we conducted a comparative analysis of feature distributions among false positives (FP), and the labeled true (T) and false (F) mutations.

**Fig 2.**
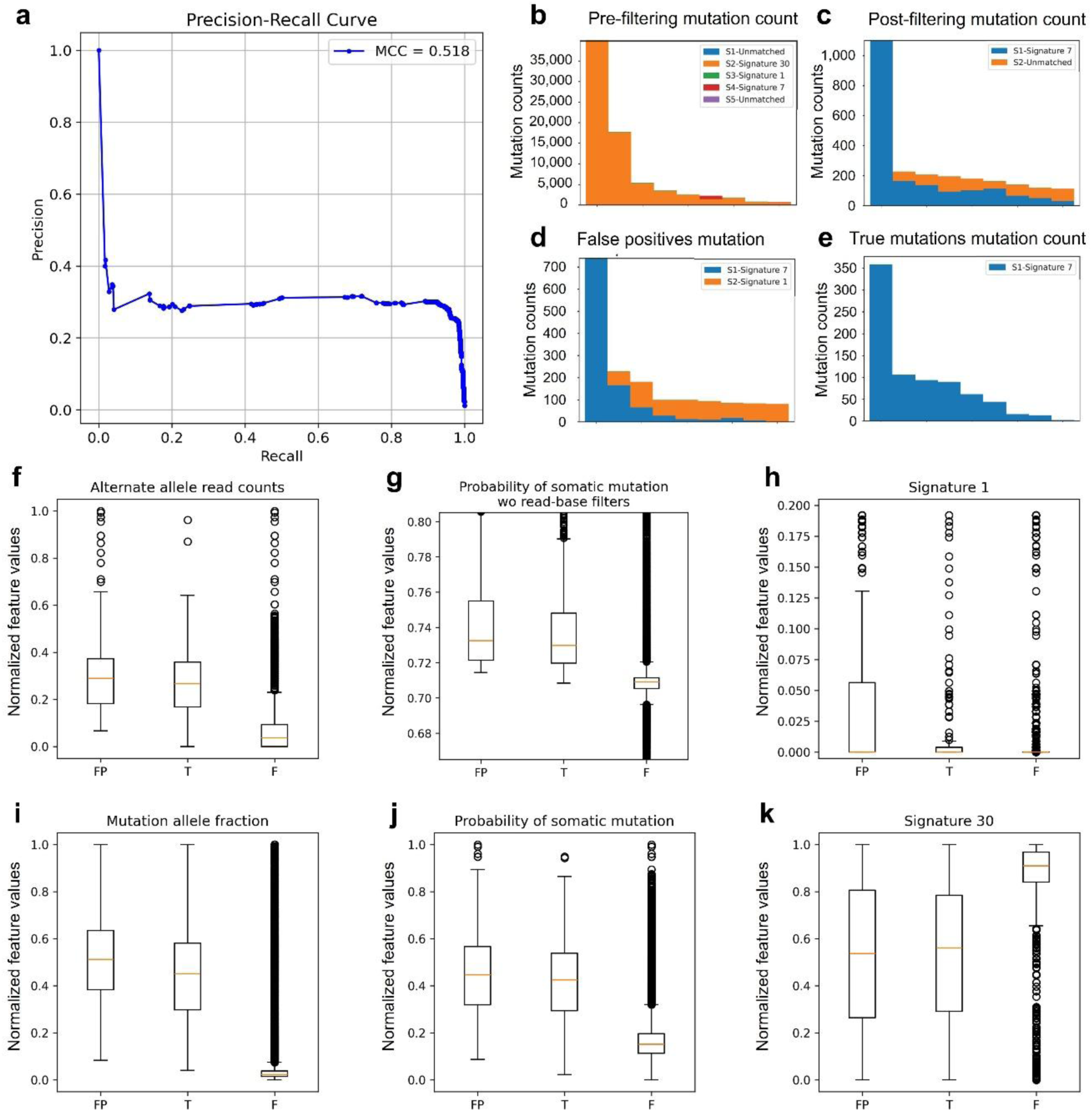
Explaining precision performance by analyzing false positive mutations. **a** Precision-Recall curve over the test set; **b-e.** Mutation count and their association with mutational signatures for pre-filtering mutations (b); post-filtering mutations (c); false positive mutations (d); true mutations (e). **f-k.** Comparing normalized feature values between FP, T and F mutations over the numerical features: **f.** Alternative allele read counts; **g.** Probability of somatic mutation without read-base filters; **h.** Mutation signature 1; **i.** Mutation allele fraction; **J.** Probability of somatic mutation; **K.** Mutation signature 30.

First, we applied SignaturAnalyzer^13–15^ on the test set before and after removing FFPE noise (Fig. 2b-c). Before removing the FFPE noise, the vast majority of the mutations are associated with signature 30 which is characterized by C>T mutation (Fig. 2b). In contrast, following noise filtering the majority of the mutations are correlated with signature 7, which is a known UV signature associated with melanoma (Fig. 2c). Similarly, we analyzed FP mutations which are potentially noise mutations that FFixR classified as true mutations. Remarkably, these mutations are associated with signature 7 (Fig. 2d), as do the true labeled mutations (Fig. 2e). Moreover, another subset of these mutations is associated with signature 1 (Fig. 2d), which is associated with aging and is commonly present in many cancer types. We next analyzed the distribution of additional features, comparing the FP mutations with true and false mutations, as originally labeled by the DNA data. We found that the FP mutations are significantly more similar in terms of their values to the true mutations than to the false mutations in every feature. In some cases, the FP mutations are even more separated from the false mutations as compared to the true mutations (Fig. 2f-k).

Overall, these results suggest that many of the mutations initially labeled as noise but predicted as true by FFixR, are likely to be genuine somatic mutations. While our baseline assumption is that all mutations detected in the RNA should also be present in the DNA, it is important to note that the DNA and RNA were not co-isolated from the exact same tissue section. Given the inherent spatial heterogeneity of tumors, it is plausible that some true somatic mutations are captured in the RNA but absent from the DNA sample, leading to their misclassification as noise. The feature-based comparison between mutation groups presented here supports this explanation and implies that the actual performance of our model is likely underestimated.

### SHAP value investigation reveals feature importance and interactions

To interpret the model’s prediction, we used SHAP (SHapley Additive exPlanations) values which quantify the contribution of each feature to the model’s output (Methods). This analysis highlights the strong predictive power of *mutation allele fraction*, along with *alternate allele count* and Mutect’s computed *Probability of somatic mutation without read-base filters* features, as the most dominant features (Fig. 3a). Indeed, it is expected that artefacts will have lower allele frequency and alternate read count, making their distinction from low clonality mutations challenging. In contrast, Mutect’s probability demonstrates a more complex behavior that can be explained in combination with additional features, as detailed below. Finally, we found that the assignment probability of a mutation to Signature 30 is also important and negatively correlates with true mutations, as depicted in Fig. 3a.

**Fig 3.**
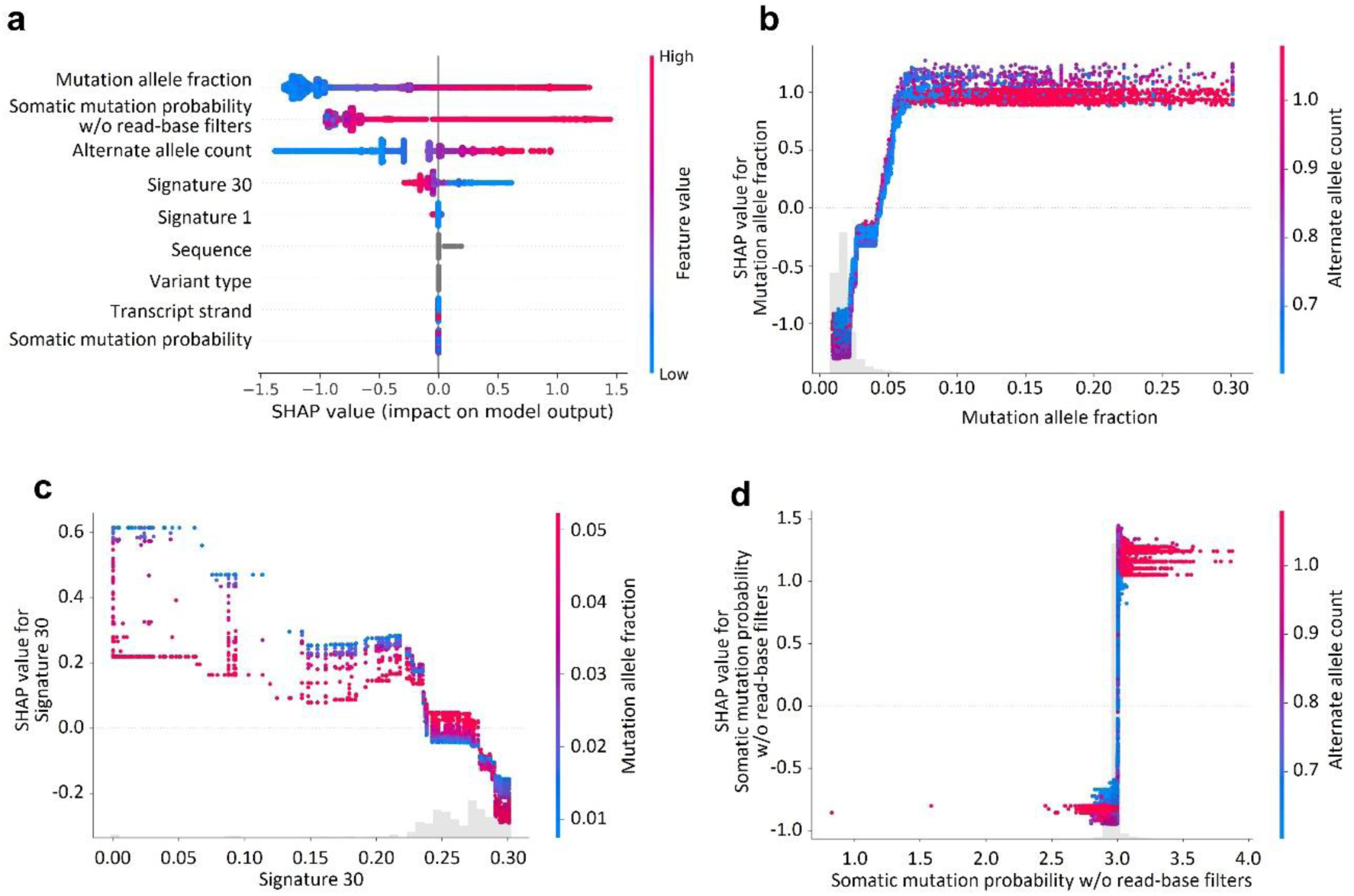
SHAP analysis and feature interaction. **a** SHAP value summary plot depicting the impact of each of the features on model output. Features’ values (shown by coloring) as a function of SHAP value (x-axis) showing the relation between each feature’s value and model’s prediction; **b-d.** Interaction plots illustrating the SHAP value dependencies between specific feature pairs. Values of the first feature shown as x-axis position, and the values of the interacting feature shown as coloring; SHAP value shown as y-axis position. A light gray histogram along the bottom axis illustrates the distribution of the x-axis feature values across the dataset: **b** Interaction between Mutation allele fraction and Alternate allele count. **c** Interaction between Signature 30 and Mutation allele fraction. **d** Interaction between Somatic mutations probability w/o read-base filter and Alternate allele count.

To further advance model explainability we performed feature interaction analysis (Methods). Considering the interaction between the alternate allele read counts and the mutation’s allele fraction, we first observed a rapid increase in SHAP value as the mutation allele fraction increases up to ∼0.2, after which the SHAP value plateaus. This suggests that small increases of this feature in the low range have a large positive impact on the model prediction, but this influence saturates beyond a certain point. As expected, the model identifies a positive interaction between mutation allele fraction and alternate allele read counts, such that it gives more weight to high allele fractions if they are also supported by a high number of reads. In addition, we found that the model recognizes that a large number of alternate reads, even at low allele fraction, can still indicate a reliable variant, for instance, in high-coverage regions or subclonal events. This suggests that alternate allele count is partially compensating for a low fraction in the model’s logic (Fig. 3b).

Another intriguing example is the interaction between probability of association with signature 30 and mutation allele fraction (Fig. 3c). Within an intermediate Signature 30 range (0.24–0.28), the model appears relatively insensitive to Signature 30, allowing allele frequency to drive the prediction more directly. Outside this range, the model’s behavior shifts. At high Signature 30 values (>0.28), an enrichment for noise-associated mutations is observed, and at low Signature 30 values (<0.24), an enrichment for true mutations is found. In both of these ranges, one might expect that higher allele fraction values would, as in the intermediate Signature 30 range, shift the model’s prediction toward a true mutation classification. However, the model demonstrates an inverse response in these regions. This counterintuitive behavior is explained by the distribution of the feature *Probability of somatic mutation without read-base filters*. Specifically, within both the high and low Signature 30 intervals, instances with low allele fraction values are associated with disproportionately high values of this probability feature, while cases with high allele fraction values exhibit comparatively low probability values. This pattern strongly influences the model’s output, overriding the expected effect of allele fraction alone (Supp Fig 3).

Finally, the feature *probability of somatic mutation without read-base filters* exhibits a step-like effect. Specifically, values below 3 are associated predominantly with noise mutations, whereas values above 3 are strongly correlated with true somatic mutations. Notably, as shown in Fig. 3d, the transition point around a value of 3 is characterized exclusively by low alternate allele read counts, while values outside this transition zone are mostly associated with high alternate allele read counts.

Together, these interaction patterns highlight the model’s nuanced decision-making, in which both individual feature values and their contextual combinations influence prediction. By capturing non-linear and compensatory relationships, the model reflects a sophisticated understanding of variant reliability. This layered explainability not only reinforces the model’s interpretability but also provides biological insights into the distinguishing characteristics of true versus artefactual mutations.

### Evaluating FFixR performance on additional melanoma dataset

We next evaluated the performance of FFixR on an independent melanoma cohort by Freeman et al.^16^, which included both fresh-frozen and FFPE-preserved samples. Among the cases with matched DNA and RNA sequencing from the same biopsy, 10 samples were frozen and 5 were FFPE-preserved. To generate an initial set of putative somatic mutations, we again applied the RNA-MuTect-WMN pipeline^8,9^. It should be noted that while we assume that all RNA true mutations should also be present in the DNA, this assumption is not entirely accurate, as in this data as well, the DNA and RNA were not co-isolated from the exact same tissue fragment.

In the five FFPE-preserved samples, FFixR achieved a Matthews Correlation Coefficient (MCC) of 0.381, with a precision of 0.204 and a recall of 0.845 — lower than the performance observed in the previous cohort. Upon further examination, we found that only 3.74% of DNA-identified mutations were also detected in the RNA, indicating substantial discordance between the two samples. In the fresh-frozen samples, the overlap between RNA and DNA somatic mutations was similarly limited: 4.7% of DNA mutations were observed in RNA, and only 18% of RNA mutations were supported by the DNA data. This highlights the intrinsic heterogeneity between RNA and DNA sequencing in this dataset. For comparison, Yizhak et al.^9^ analyzed DNA and RNA from TCGA tumor samples co-isolated from the same biopsy, demonstrating that in that case approximately 35% of somatic mutations identified in DNA were also detectable in the corresponding RNA. Moreover, in the Van Allen et al. cohort, 18% of DNA mutations overlapped with RNA, suggesting that the degree of RNA–DNA divergence is more pronounced in the Freeman et al. dataset. This increased discordance likely introduces substantial mislabeling of true mutations as false positives, contributing to the apparent reduced performance of the model. This interpretation is further supported by the observation that recall remains largely stable, whereas precision exhibits a marked decline.

Given the clinical importance of Tumor Mutational Burden (TMB) as a biomarker for predicting response to immune checkpoint inhibitors (ICI) and patient prognosis, we next sought to evaluate the utility of our model in this context. TMB, defined as the total number of non-silent somatic mutations in a tumor sample, is typically estimated based on DNA sequencing data. However, prior studies, including our own^8,10^ have shown a strong linear correlation between DNA-based and RNA-based mutation counts. Notably, RNA-based TMB, which reflects only expressed mutations, has shown improved predictive power in certain contexts^8^. Leveraging this relationship, we investigated the ability of our model to recover a reliable estimate of RNA-derived TMB that aligns with the DNA-based gold standard. The ability of FFixR to restore this correlation—despite imperfections in recall and precision—would underscore its potential for clinically meaningful applications.

To evaluate this, we examined the correlation between RNA-based and DNA-based TMB across various sub-cohorts. In the FFPE-preserved samples, no significant correlation was observed prior to applying FFixR (R² = 0.014; Fig. 4a). As expected, the frozen tissue samples showed a strong linear correlation between RNA and DNA TMB values (R² = 0.842; Fig. 4b). Notably, after applying FFixR to the FFPE samples, the correlation improved substantially (R² = 0.881), indicating that the model effectively mitigated FFPE-induced noise. When combining the 15 samples (10 frozen and 5 FFPE), we observed a moderate overall correlation (R² = 0.760), slightly lower than that of each sub-cohort individually. This decrease in overall correlation may result from residual FFPE-induced artifacts in the RNA data that inflate the apparent TMB, or from overcorrection by the model that inadvertently removes true mutations, thereby lowering TMB estimates. On the one hand the model’s performance metrics, particularly its moderate precision, suggest the former explanation is more likely. On the other hand, since the frozen samples reflect the true RNA-DNA relationship, the lower slope in the FFPE set may indicate the latter.

**Fig 4.**
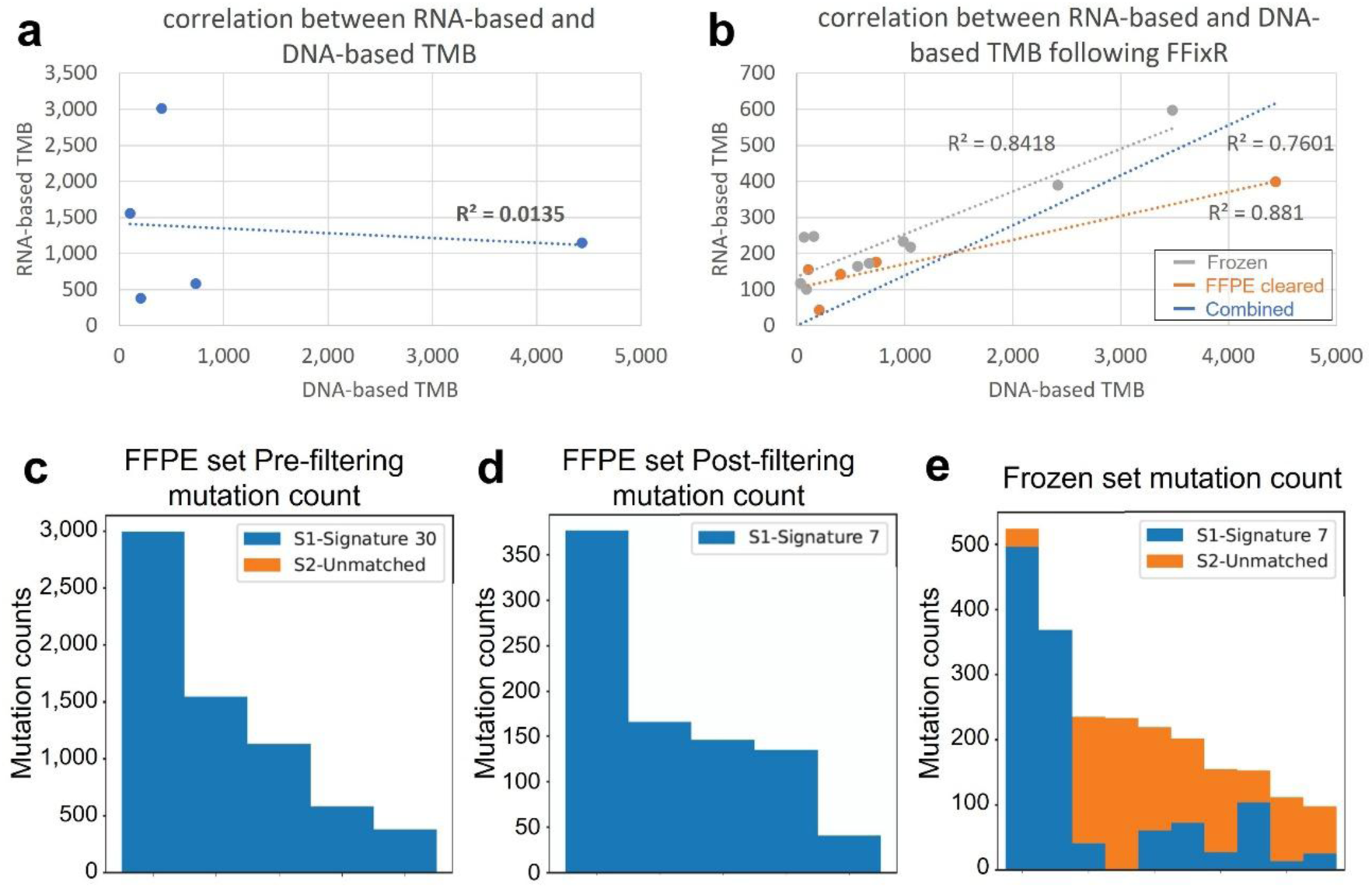
Analysis of the Freeman et al. cohort. **a** RNA-based TMB of the 5 FFPE samples before applying FFixR vs. DNA-based TMB. **b** Correlation between DNA-based TMB and the RNA-based TMB of the 10 frozen samples in gray, the 5 FFPE samples after applying FFixR in orange and both sub cohorts combined in blue. **c-e** Mutation count and their associated signatures in the 5 FFPE samples before applying FFixR (c), following FFixR application (d) and in the 10 frozen samples.

Finally, we applied SignatureAnalyzer to the various Freeman et al. sub-cohorts to assess the impact of FFixR on mutational signature composition. We found that prior to noise filtering, the five FFPE-preserved samples exhibited a predominance of mutational signature 30, which is commonly associated with FFPE-induced artefacts (Fig. 4c). Following application of FFixR, these artefactual mutations were effectively removed, and the remaining mutation set was primarily enriched for signature 7, a pattern characteristic of UV-induced damage and typical of melanoma (Fig. 4d). Similarly, the 10 frozen samples—unaffected by FFPE artefacts—also showed a dominant contribution from signature 7 (Fig. 4e), consistent with biological expectations. These findings further support the ability of FFixR to selectively remove FFPE-associated noise while preserving biologically meaningful somatic mutations.

## Discussion

In this study, we present a robust computational framework for identifying and removing FFPE-induced artifacts from RNA-seq data, addressing a critical limitation in leveraging archived FFPE samples for genomic analyses. Our approach, FFixR, implemented via an XGBoost model, effectively discriminates between true somatic mutations and FFPE-associated noise despite the constraints posed by the single-stranded nature of RNA. This distinguishes our method from traditional DNA-focused artifact detection tools that rely primarily on strand bias metrics and are inapplicable to RNA-seq data.

The results highlight the efficacy of our approach across different melanoma cohorts. In the primary dataset, we achieved high recall (∼0.92) and eliminated the overwhelming majority (∼97%) of FFPE-induced artifacts. Notably, precision remained moderate (∼0.30), especially for samples with low somatic mutation counts, suggesting that some true positives may be mislabeled due to the absence of matched DNA data for certain RNA-detected variants. The comparable feature profiles of false positives and true positives further underscore the biological and technical complexities underlying this data, implying that certain RNA variants classified as noise might indeed be legitimate somatic mutations arising from tumor heterogeneity or expression-driven dynamics.

Importantly, our results reveal the nuanced interplay between variant features, with allele fraction, alternate read count, and signature-associated probabilities exerting dominant roles in the classification process. The use of SHAP analyses and feature interaction modeling provides critical insights, highlighting thresholds and saturation points that govern the model’s behavior. For example, low variant allele fraction combined with high alternate read counts can be a strong indicator of a true variant.

The improved correlation between RNA- and DNA-derived TMB in FFPE samples following application of FFixR (R² rising from 0.014 to 0.881) demonstrates its clinical and biological value. The ability to obtain accurate RNA-based TMB estimates is pivotal for extending immunogenomic analyses to FFPE archives, facilitating the exploration of predictive biomarkers for immunotherapy across wider patient populations.

Although the approach was developed and validated in melanoma, it is not reliant on melanoma-associated mutational signatures and incorporates signatures 1 and 30 as generalized indicators of aging and FFPE-associated damage, respectively. This suggests its potential application across other malignancies and tissue types. Nonetheless, further studies using larger, multi-tissue FFPE collections and matched DNA-RNA data will be necessary to rigorously assess its generalizability and clinical utility.

In conclusion, the presented framework allows for the accurate recovery of somatic mutations from FFPE RNA-seq data, supporting its use in both clinical diagnostic and research settings. By addressing a long-standing barrier in FFPE data utilization, this approach expands the potential for transcriptomic studies to guide precision medicine and deepen our understanding of the molecular underpinnings of cancer.

## Methods

### Detection of somatic mutations in RNA-seq data

Raw RNA-seq data was processed using the **R**NA_MuTect_WMN pipeline^8^, which enables detection of somatic mutations from RNA sequencing data in the absence of matched-normal samples. This pipeline produced MAF files containing candidate somatic mutations and FFPE-induced artefacts, while excluding germline variants.

### Extracting mutational signature cosine similarity

The second step of our pipeline involved applying the SignatureAnalyzer Python package to the RNA-derived MAF files. This tool assigns each mutation a score between 0 and 1, representing its probability of association with known COSMIC mutational signatures. For model development, we selected signatures that correlated with FFPE artifacts^11^, such as signature 1 that is commonly associated with aging, and signature 30, which has been reported in a subset of breast cancers^14^.

### Data preparation and model training

We trained our model using RNA-seq data from 39 melanoma patients^12^. As the ground truth we used the set of mutations derived from the matched whole-exome sequencing (WES) data, as reported in the original publication. As an independent test set, we randomly set aside 9 out of the 39 samples using np.random.seed(42) for reproducibility. The remaining 30 samples were used to train the model using the XGBoost Python package in a 5-fold cross-validation manner. In each fold, 24 samples (80% of the training set) were used for training, and the remaining 6 (20%) served as the fold-specific validation set (see Supp. Table 2). All variants were aggregated into a single DataFrame per fold, resulting in varying numbers of mutations across folds due to differences in sample sizes.

To optimize model performance, we performed hyperparameter tuning to maximize the average Matthews Correlation Coefficient (MCC) across folds, a metric particularly suited for imbalanced classification tasks. Precision and recall were also used as secondary evaluation metrics. The final set of hyperparameters is as follows is available in supp. Table 3. All other hyperparameters were left at their default values. Finally, to predict mutation labels in the validation set, we applied majority voting across the five models trained in the cross-validation folds.

### Feature extraction and processing

The model was trained using a set of features that characterize the frequency and count of the mutant versus reference alleles, the likelihood ratio of a candidate mutation being true versus a sequencing artifact, the tri-nucleotide sequence context, cosine similarity to known COSMIC mutational signatures (specifically Signatures 1 and 30), and strand orientation (Supp Table 1).

The sequence context was encoded as a 4-character string representing the base immediately upstream, the reference base, the alternate base, and the downstream base. Considering the complementary strand, this encodes for 96 possible sequences^14^. Variant type was one-hot encoded for one of the 13 possible types that were present in the data we worked with (see Supp Table 1). Strand orientation was encoded as a Boolean variable. All other features were numerical and transformed using the formula:

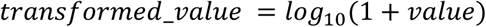

In the specific case of the feature *Probability of somatic mutation without read-base filters*, where the values assigned during MuTect process range between ∼-19,000 and ∼9,000, we first snapped all values at −1,000, such that values smaller than −1000 were converted to −1,000. This process affected 0.05% of the mutations. Next, we added +1,000 before applying the *log_10_(1 + value)*. This transformation process was implemented in order to bring all features to be positive (see Supp Figs 1,2, Supp Table 1).

### Analyzing mutational signatures

The SignatureAnalyzer Python package was used to compute the cosine similarity between each mutation and known COSMIC mutational signatures, as described above. In addition to generating these per-mutation similarity scores, the package was also employed to visualize mutational signature distributions across various cohorts. These included, for example, the validation set prior to the FFPE noise-filtering step, as well as subsets of mutations classified as false positives following noise removal.

### SHAP features importance and features’ interaction analysis

SHAP (SHapley Additive exPlanations) is a widely used method for interpreting machine learning models by quantifying the contribution of each feature to individual predictions. We employed the SHAP Python package to analyze feature importance within our model. The resulting SHAP summary plots illustrate the impact of each feature on the model’s output across all mutations, capturing both the predictive strength and the direction of the association—i.e., whether higher feature values are associated with classification as true mutations (class 1, indicated in red) or noise (class 0, indicated in blue). To investigate feature interactions we employed the following function:

shap.plots.scatter(explanation[:, feature], color=explanation)

This function visualizes the effect of individual features. This command was applied iteratively across all features; however, only those with the highest overall importance are presented (Fig. 3).

### Freeman et al. cohort analysis

Our analysis pipeline was applied to the entire dataset, which included two sub-cohorts: five FFPE-preserved samples and 10 frozen samples. All 15 samples were first processed using the RNA_MuTect_WMN pipeline to identify putative somatic mutations from RNA-seq data without matched-normal samples. Subsequently, only the FFPE samples were further processed using FFixR to remove FFPE-induced artifacts. TMB was estimated for each sample by quantifying the number of non-silent mutations reported in the respective MAF files for both DNA and RNA. Similarly, for the FFPE samples after FFixR implementation the TMB was calculated from the set of mutations remaining after filtering.

## Data availability

RNA sequencing data from 39 melanoma patients of the Van Allen et al. dataset^12^ is available online in dbGAP database under accession number phs000452.v3.p1. List of mutations in the DNA for the same cohort is available in the paper^12^ as tables1.mutation_list_all_patients in the Supplementary Material section. RNA sequencing data from 15 melanoma patients of the Freeman et al. dataset^16^ (MGH cohort) is available in dbGAP with accession number phs002683.v1.p1.

## Code availability

The methods described in this manuscript have been implemented in a tool called FFixR, which is freely available at the GitHub repository. The repository also includes a readme file describing the inputs, outputs and entire pipeline: https://github.com/yizhak-lab-ccg/FFixR.

## Author contributions

K.Y. supervised. K.Y. and O.L. jointly wrote the main manuscript text. O.L. prepared the figures and performed the analysis. Both authors reviewed and approved the final manuscript.

## Competing interests

The authors declare no competing interests.

**Correspondence** and requests for materials should be addressed to Keren Yizhak.

## Supporting information

Supplementary figure 1 Numerical features raw values

Supplementary figure 2 Numerical features processed values

Supplementary figure 3 Probability of somatic mutation wo read-base filters in high-low regions of Signature 30 and Mutational allele fraction

Supplementary table 1 features

Supplementary table 2 Training and test set samples

Supplementary table 3 Models hyperparameters

## Supplementary information

Supplementary table 1 - Features’ table

Supplementary table 2 - Training and test set samples Supplementary table 3 - Model’s hyperparameters Supplementary Fig. 1 - Numerical features raw values Supplementary Fig. 2 - Numerical features processed values

Supplementary Fig. 3 - Values of Probability of somatic mutation without read-base filters for the various combinations of High/ Low values of Signature 30 and Mutation allele fraction features

## Abbreviations

FFPE: Formalin-fixed paraffin-embedded
RNA-seq: RNA sequencing
DNA-seq: DNA sequencing
MAF: Mutation Annotation Format
ML: Machine Learning
MCC: Matthews Correlation Coefficient
RNA-MuTect-WMN: RNA-MuTect Without Matched Normal
FP: False Positive mutations
T: True labeled mutations
F: False labeled mutations
SHAP: SHapley Additive exPlanations
TMB: Tumor Mutational Burden
WES: Whole-Exome Sequencing

## Notes

https://dbgap.ncbi.nlm.nih.gov/beta/study/phs000452.v3.p1/

https://dbgap.ncbi.nlm.nih.gov/beta/study/phs002683.v1.p1/

### Competing Interest Statement

The authors have declared no competing interest.

