## Supplementary figures and images for "FFixR: A Machine Learning Framework for Accurate Somatic Mutation Calling from FFPE RNA-Seq Data in Cancer"

### Supplementary figure 1 Numerical features raw values

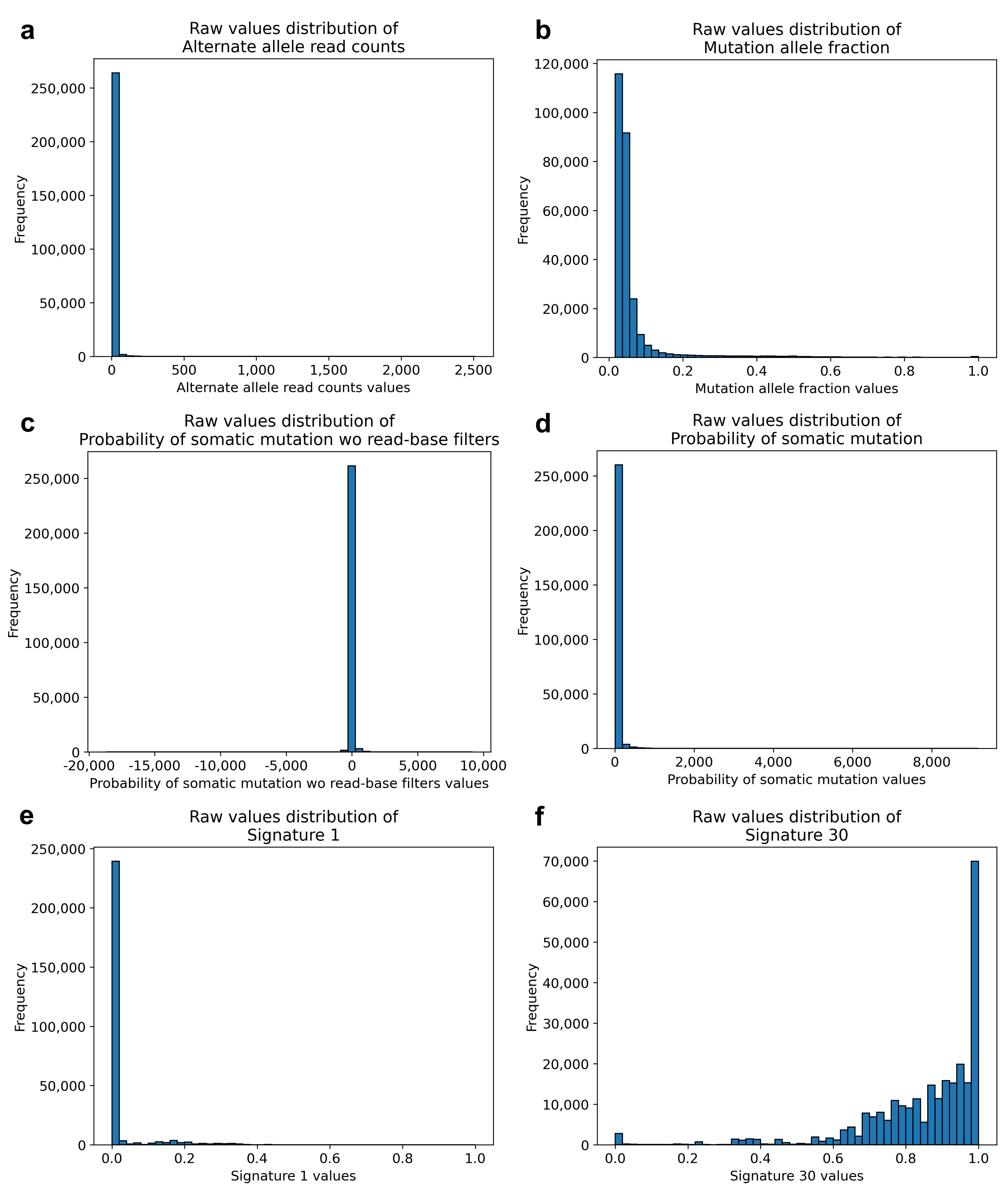

### Supplementary figure 2 Numerical features processed values

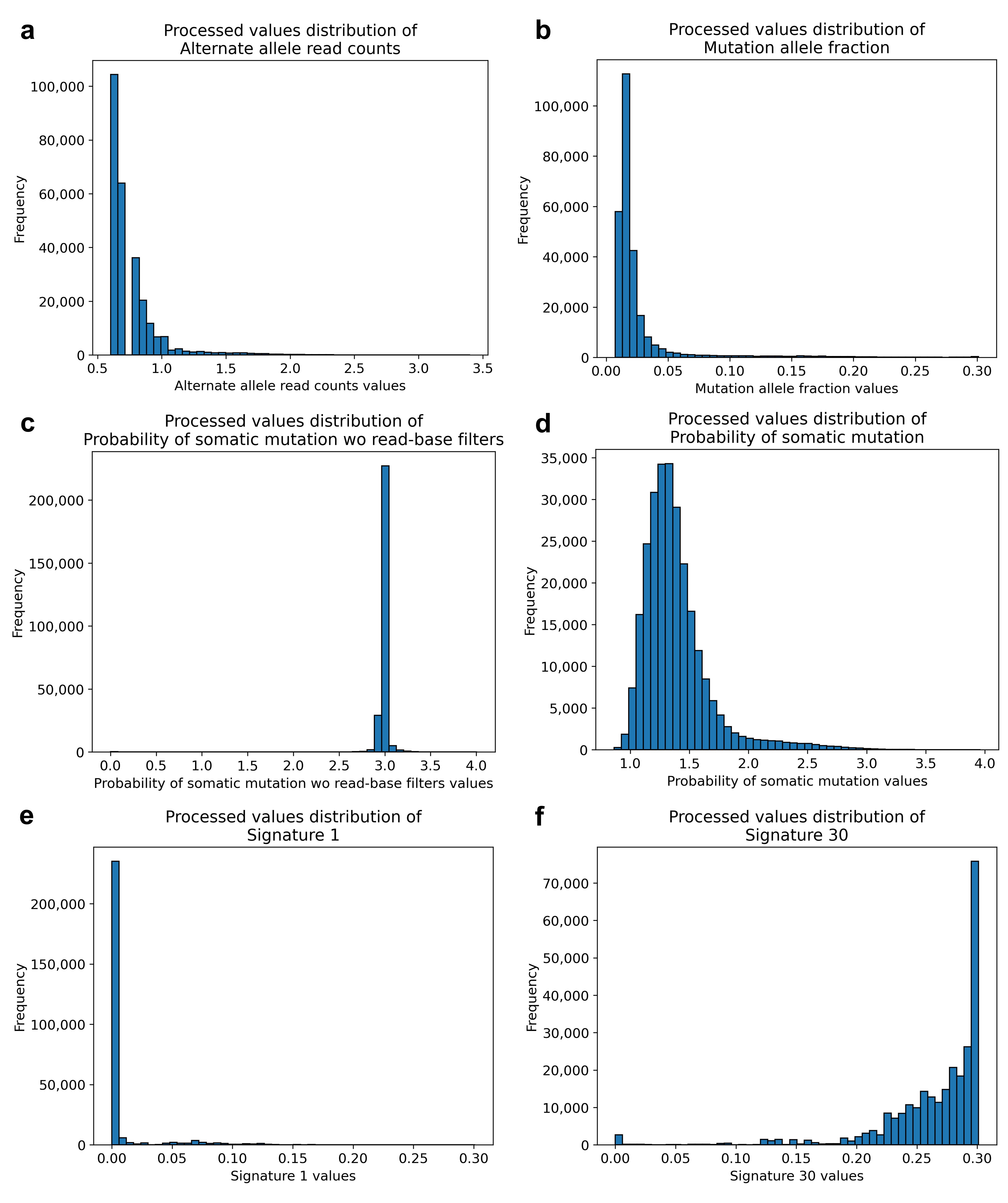

### Supplementary figure 3 Probability of somatic mutation wo read-base filters in high-low regions of Signature 30 and Mutational allele fraction

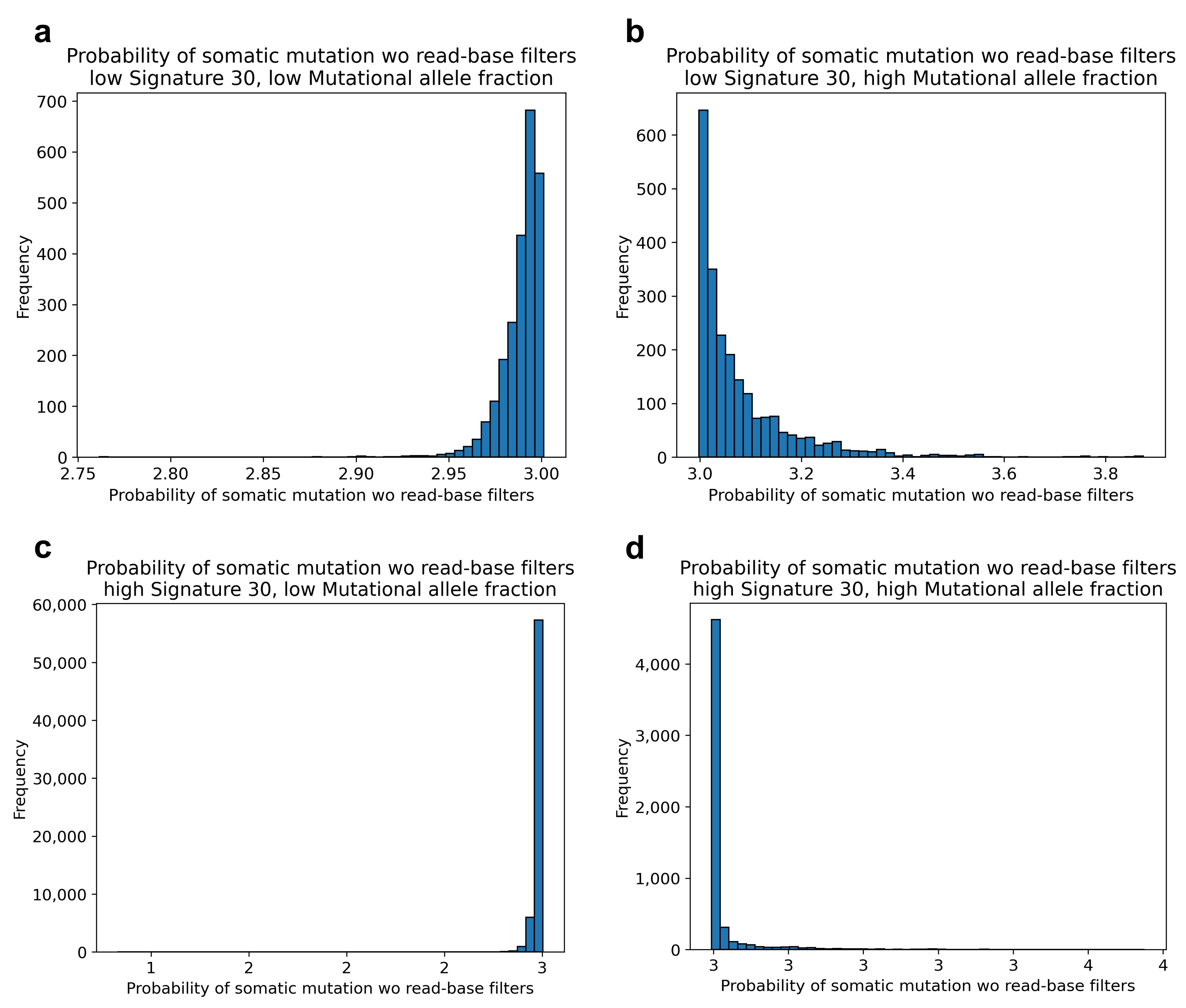
